# The genetics of assisted gene flow: immediate costs and long-term benefits

**DOI:** 10.1101/2021.04.20.440707

**Authors:** Jared A. Grummer, Tom R. Booker, Remi Matthey-Doret, Pirmin Nietlisbach, Andréa T. Thomaz, Michael C. Whitlock

## Abstract

Plant and animal populations are facing several novel risks such as human-mediated habitat fragmentation and climate change that threaten their long-term productivity and persistence. With the genetic health of many populations deteriorating due to climate change outpacing physiological adaptation, human interventions in the form of assisted gene flow (AGF) may provide genetic variation to adapt populations to predicted climate change scenarios and result in more robust and productive populations. We ran genetic simulations to mimic a variety of AGF scenarios and measured their outcomes on population-level fitness to answer the question: in which circumstances is it worthwhile to perform AGF? Based on the parameters we explored, AGF may be harmful in certain situations over the short term (e.g., the first ∼10-20 generations), due to outbreeding depression and introducing deleterious genetic variation. Moreover, under many parameter sets, the benefits of AGF were relatively weak or took many generations to accrue. In general, when the adaptive trait is controlled by many loci of small effect, the benefits of assisted gene flow take much longer to realize–potentially too long for most climate-related management decisions. We also show that when translocation effort is divided across several generations and outbreeding depression is strong, the recipient population experiences a smaller decrease in fitness as compared to moving all individuals in a single effort. Importantly, in most cases, we show that the genomic integrity of the recipient population remains relatively intact following AGF; the amount of genetic material from the donor population typically ends up constituting no more of the recipient population’s genome than the fraction introduced. Our results will be useful for conservation practitioners and silviculturists, for instance, aiming to intervene and adaptively manage so that populations maintain a robust genetic health and maintain productivity into the future given anthropogenic climate change.

## INTRODUCTION

Natural populations are currently facing a multitude of anthropogenic threats, such as climate change and habitat loss, alteration, and fragmentation, that lead to fitness reductions and population declines (Klenner and Arsenault, 2009; Pacifici et al. 2017). In many species, a once suitable environment becomes of lower quality and the local population is either forced to migrate, adapt, exhibit phenotypic plasticity, or become extinct (Hamilton and Miller 2016). Many populations cannot easily adjust to climate change because migration to suitable habitat is not possible due to habitat fragmentation and the lack of suitable habitat corridors (Hoffman and Sgrò 2011). Phenotypic plasticity may provide temporary relief (Levis and Pfennig 2016). However, rapid anthropogenically-based climate change is outpacing the natural process of environmental adaptation through natural selection (Gonzalez et al. 2013).

Natural selection allows populations to adapt to environmental conditions, but adaptation requires genetic variation that may be lacking in certain populations. Particularly when environmental conditions are changing rapidly, populations may experience a lag in adaptation to the environmental conditions they experience, and thereby suffer fitness reductions (e.g., Browne et al. 2019). Recently, human activities have caused the climate to change particularly fast (Stocker et al. 2014). As a consequence of changing climates and environments, an increasing number of populations experience an evolutionary lag and struggle with the pace of environmental change (e.g., Uecker et al. 2014; Wilczek et al. 2014; Radchuk et al. 2019; Klausmeier et al. 2020).

Many commercially relevant species such as crop plants and trees for timber production have been studied for their capacity to adapt to changing climates (Anderson 2016). For instance, Wang et al. (2010) showed that greater harvests of lodgepole pine (*Pinus contorta*), a commercially harvested tree species in British Columbia, Canada, could be achieved when planting efforts account for genetic and predicted climatic effects. Similarly, many marine organisms of economic importance for harvesting or ecotourism, such as coral reefs, are also under threat due to ocean acidification and rising temperatures resulting from acute and severe episodic ocean-climate events (Baker et al. 2008). In some systems, natural populations show differential responses to bleaching (Hughes et al. 2003), suggesting that gene flow from other populations could be beneficial for those populations suffering more severely from bleaching events. Indeed, genomic data and biophysical modeling for a coral in the Great Barrier Reef suggest that gene flow from lower to higher latitude populations can provide beneficial heat-tolerant alleles as the climate warms (Matz et al. 2018), in spite of moderate levels of selection against migrants in other coral systems (Kenkel et al. 2015). In terrestrial populations, gene flow from populations in drier regions could help populations in more mesic regions that are predicted to experience a higher frequency of droughts via climate change (e.g., Exposito-Alonso et al. 2018). However, natural gene flow may not always be possible because of natural dispersal barriers or anthropogenic habitat fragmentation. Over recent decades, habitat fragmentation has increased considerably, both in terrestrial (Haddad et al. 2015) and aquatic (Grill et al. 2019) habitats, likely often inhibiting the natural process of adaptation via gene flow.

Assisted gene flow (AGF), i.e., human-mediated translocation of individuals from other populations to pre-adapt a population to a changing environment, has been proposed as a way to introduce genetic variation into populations experiencing fitness declines via maladaptation (Aitken and Whitlock 2013; Uecker et al. 2014; Tomasini and Peischl 2020). AGF is a specific type of assisted migration. Assisted migration includes translocations both within and outside of species’ ranges but is typically focused on movement of individuals outside of current species’ ranges, whereas AGF refers to translocations among existing populations. Although related concepts, a distinction should be made between AGF, genetic rescue, and evolutionary rescue. Genetic rescue is an increase in the fitness of small populations owing to the immigration of new alleles (Tallmon et al. 2004), and is generally considered to occur when population fitness increases by more than what can be attributed to the demographic contribution of immigrants (Ingvarsson 2001). Thereby, genetic rescue alleviates the deleterious consequences of inbreeding in small populations. Evolutionary rescue occurs when a population adapts, through natural selection, to a changing environment and results in demographic stabilization, population persistence and rescue from extinction (Bell 2013). Thus, evolutionary rescue is typically invoked in large populations and includes adaptation to a changing environment, whereas genetic rescue occurs in small populations by reducing inbreeding depression and promoting heterosis.

While the intended consequences of genetic or evolutionary rescue are to prevent a population from going extinct, AGF aims to prevent extinction of threatened species or to promote productivity in a species of economic importance (e.g., Aitken and Bemmels 2016). AGF is typically invoked as a management action when a population is no longer adapted to its environment. To achieve this goal, individuals harboring pre-adaptive alleles (alleles that cause local adaptation to a particular environmental trait) are translocated to rescue an imperiled population (Aitken and Whitlock 2013). Even if their environment has been stable, small populations that suffer from inbreeding depression and fixed deleterious genetic variation may benefit from genetic rescue via AGF (Gaggiotti and Hanski 2004).

The introduction of genotypes from foreign sources poses its own risks. For instance, genetic swamping resulting from “hybridization” between distinct populations can lead to the loss of a population’s genetic integrity and potentially genomic extinction (Todesco et al. 2016).

Additionally, individuals from foreign populations may carry alleles which are maladaptive under the local conditions or that may create outbreeding depression in combination with local alleles. Locally deleterious variation could be responsible for adaptation to environmental dimensions other than the ones of interest (e.g., Etterson and Shaw 2001). For example, AGF may be planned with the aim of aiding adaptation to warming temperatures, but the source or “donor” population may be adapted to a different seasonal cycle and thus exhibit a phenology that is incompatible with the conditions at the new location. Similar differences between the environments of the source and recipient populations may exist in precipitation, exposure to the weather, or other circadian and circannual (day length, seasonality) traits. Furthermore, although populations may be selected for their ability to handle warmer temperatures, light limitations from lower latitude populations may restrict their use in poleward assisted migrations (Wadgymar et al. 2015; Huffeldt 2020). Finally, because alleles sometimes interact poorly with alleles from other populations (outbreeding depression), some alleles that work well in one genetic context can cause fitness declines in a novel genetic context (Templeton et al. 1986, Frankham et al. 2011). Such locally deleterious alleles will initially be in strong linkage disequilibrium with the introduced beneficial alleles, making them even more potentially consequential. For reasons such as this, it is important to take into consideration the level of population relatedness and environmental differences between donor and recipient populations, but it is currently unclear how these negative effects can reverse or ameliorate the benefits of AGF.

Conservation managers have several practical decisions to make when considering whether to invoke assisted gene flow or not:

> *Migration effort*―how many individuals to translocate? Translocating a high number of individuals to increase the odds of allele frequency change in the recipient population could come at a high financial cost. Moreover, the resulting high proportion of foreign individuals may dilute or even completely replace the “native” composition of the recipient gene pool. Conversely, adding too few individuals might not introduce the pre-adaptive alleles at a high enough frequency to remain in the population.
>
> *Translocation strategy*―translocate all individuals at once, or over several generations (a “pulsed” strategy)? Will a pulsed translocation effort spread across several generations ease the transition of translocated individuals into a foreign ecosystem?
>
> *Fitness reductions*―does the long-term gain outweigh the short-term loss? Outbreeding depression and maladaptive alleles can lead to a sharp reduction in population-level fitness following translocation; is this “fitness valley” so deep that it can impair fitness recovery and potentially lead to local extinction, or shallow enough to be transcended and ultimately lead to long-term improvements in fitness?

In this study, we investigated under which genetic circumstances it is worthwhile to perform assisted gene flow using forward-in-time individual-based simulations. We assessed the combined effect of alleles that are pre-adapted to a changing climatic variable (i.e., lead to local adaptation), maladaptive alleles (i.e., alleles that are fixed in the donor population but deleterious in the recipient population), and alleles that cause outbreeding depression through the breakdown of epistatic interactions. We simulated various genetic architectures of these traits that have positive (“pre-adaptation”) and negative (maladaptation, outbreeding depression) consequences on the fitness of the recipient population. We then tracked population mean fitness over time to determine how different genetic architectures affected the short- and long-term fitness of the recipient population. We end by discussing the implications of these results for natural resource managers and conservation practitioners.

## MATERIALS & METHODS

We conducted population genetic simulations using SimBit v4.9.30 (Matthey-Doret 2020). A glossary of terms and default parameter values for variables are listed in Table 1. We simulated the translocation of individuals from a donor population into a recipient population of either 1,000 or 10,000 diploid individuals. Individuals from the donor population were fixed for alleles that were pre-adaptive or maladaptive in the recipient population (see below). We modelled a translocation fraction (*T*_*f*_) of either 5% or 50% (simulation results of *T*_*f*_ = 0.5% are available in the Supplement). Although a translocation fraction of 50% may be unrealistic in certain scenarios, it serves as an extreme example to help visualize trends of the impact of translocation effort on fitness and the maintenance of local genetic identity. We modelled selection on fecundity so the first round of selection in all simulations occurred at the time that generation 1 was produced. Individuals were translocated in a single pulse or in five pulses separated by either 1, 2, or 4 generations, each pulse representing 20% of the total number of individuals translocated. After the introduction, the simulated recipient population evolved for 100 generations.

**Table 1.**
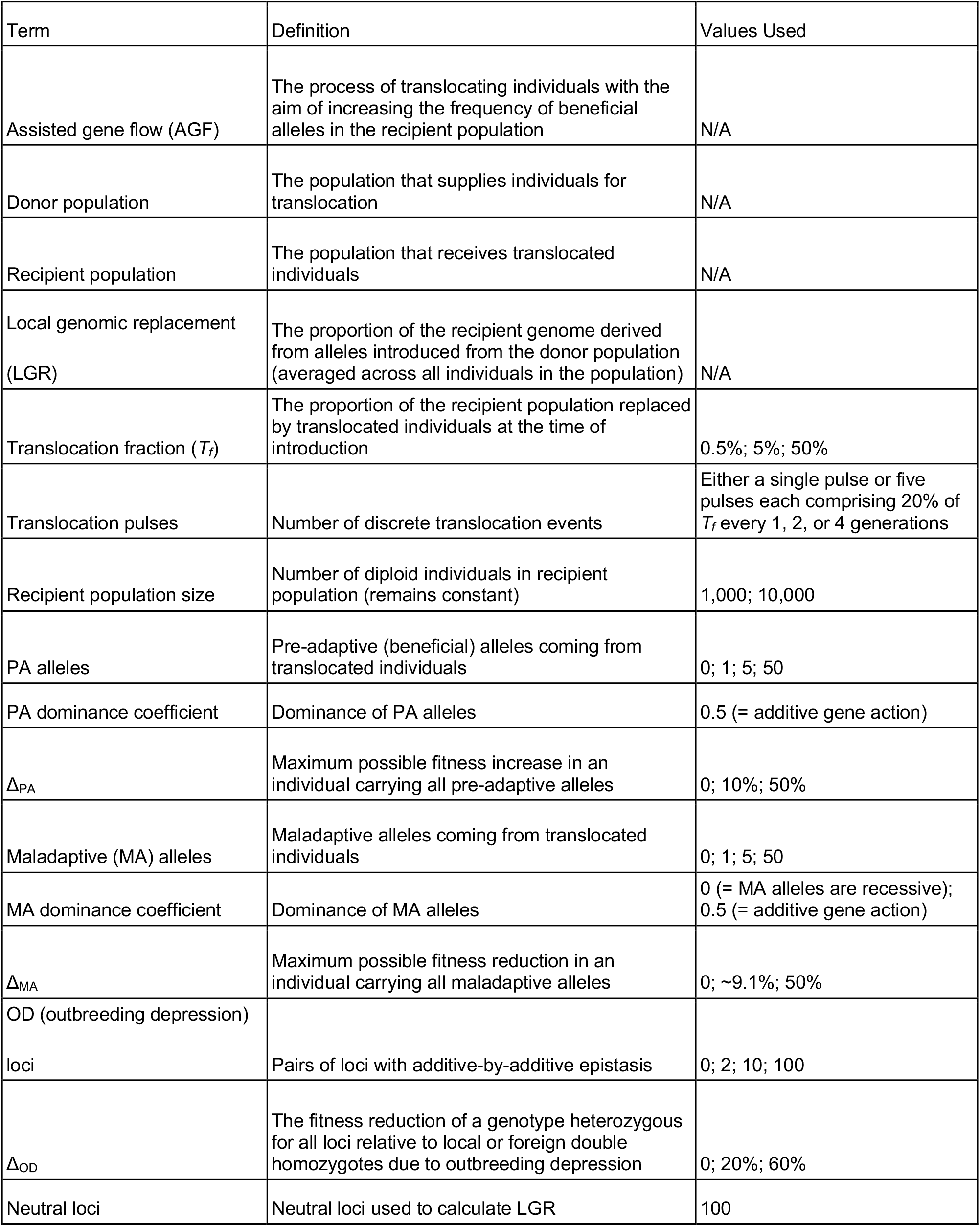
A glossary of terms and the parameters used in this study. For simulation parameters, we include the values used in our factorial simulation analysis.

Our simulations modelled alleles that were both locally pre-adapted and maladapted to the climate experienced by the recipient population. We modelled alleles that were selectively favored in the donor population (and therefore fixed there). Some of these alleles would be adaptive in the new environment of the recipient population (“pre-adapted alleles”) and other alleles have low fitness in that new environment (“maladaptive alleles”). We assumed that these alleles were absent in the recipient population until they were introduced by AGF. We parameterized the selective effects of pre-adaptive and maladaptive alleles by focusing on the overall effect of 0, 1, 5 or 50 selected loci. If an individual in the recipient population was homozygous for all pre-adaptive alleles, its relative fitness would be increased by Δ_*PA*_. Similarly, if an individual in the recipient population was homozygous for all maladaptive alleles, its relative fitness would be decreased by Δ_*MA*_. We simulated cases with Δ_*PA*_ of 10% or 50% and Δ_MA_ = ∼9.1% and 50%. Note that these Δ_*PA*_ and Δ_*MA*_ values represent the *maximum* change in fitness, e.g., when an individual is homozygous for all pre-adaptive or maladaptive alleles. As a consequence, for a given Δ_*PA*_ or Δ_*MA*_, the higher the number of loci, the lower the selection coefficient per locus. The values were chosen so that Δ_*PA*_ of 10% will compensate Δ_*MA*_ ≈ 9.1% in individuals homozygous for either all PA or MA alleles. The dominance coefficient was set to 0.5 for PA loci and 0.5 for MA loci (0 dominance for MA loci is shown in the Supplementary Material). Outbreeding depression poses an additional genetic risk of translocating individuals from other populations (Frankham et al. 2011). For example, epistatic interactions among pairs of loci may considerably reduce the fitness of some double homozygotes (Orr 1995, 1996). At equilibrium, these strongly deleterious double homozygotes rarely occur in a population. However, they may occur more commonly in individuals with mixed ancestry from differentiated populations.

Outbreeding depression (OD) was modelled by simulating pairs of loci with additive-by-additive epistasis. An individual homozygous for either foreign or local alleles at both loci in a pair had fitness of 1 + *s*_OD_ relative to the double heterozygote (double heterozygote fitness = 1). Individuals homozygous for local alleles at one locus and foreign alleles at the other had a fitness of 1 – *s*_OD_ relative to the double heterozygote. All other genotype combinations (i.e., at least one heterozygote in a pair) had a fitness of 1.0. We modelled either 0, 2, 10, or 100 pairs of such epistatic OD loci.

Following Aitken & Whitlock (2013), we parameterized outbreeding depression by focusing on the overall fitness reduction across all pairs of loci. Individuals heterozygous for all epistatic pairs had a fitness reduction of Δ_*OD*_ relative to the fittest homozygotes. We simulated cases where outbreeding depression resulted in fitness reductions of either Δ_*OD*_ *=* 20% or 60% in heterozygotes relative to the fittest double homozygotes. By way of illustration, when simulating Δ_*OD*_ = 20% across 2 pairs of epistatic loci, *s*_OD_ ≈ 0.1180, for 10 pairs *s*_OD_ ≈ 0.0226, and for 100 pairs *s*_OD_ = 0.0022. A fitness reduction of 60% resulted from 2 pairs of epistatic loci with *s*_OD_ ≈ 0.5811, 10 pairs with *s*_OD_ ≈ 0.0960, or 100 pairs with *s*_OD_ ≈ 0.0092. Though likely not indicative of intraspecific population pairings, a Δ_*OD*_ of 60% illustrates an extreme case to visualize the effect of outbreeding depression on fitness.

For the simulation results with PA and MA loci presented in the main text, we assumed multiplicative fitness effects among loci and additive interactions between alleles within loci. Fitness effects of OD loci pairs also interact multiplicatively with all other pairs and the PA and MA loci. All loci were randomly distributed onto 10 chromosomes of 10 centimorgans each with a uniform recombination rate. Thus, pairs of loci causing outbreeding depression were randomly distributed in the genome and therefore often not closely linked.

Local genomic replacement (LGR) may occur as introduced haplotypes spread to fixation in the recipient population, replacing local genetic diversity. Given the concern that an increase in local fitness may be due to a proportionate change in non-local ancestry (e.g., Harris et al. 2019), we measured local genomic replacement by including 100 neutral loci in all simulations that were reciprocally fixed in the source and recipient populations; these were placed at random along the genome. At every generation of a simulation run, we calculated the mean allele frequency across all the neutral markers to obtain a measure of the average proportion of alleles in the local population that derived from alleles introduced during AGF. Our measure of LGR therefore varies from 0.0 to 1.0, where 1.0 indicates that all neutral alleles in the recipient population are derived from the donor population.

For each combination of all parameter values summarized in Table 1, we performed 50 simulation replicates on a total of 1,536 unique parameter combinations. We measured relative fitness in the recipient population and calculated the mean fitness across replicates. We used the mean frequency of neutral alleles as a measure of local genomic replacement.

### Data availability statement

Code to perform and analyze the simulations as well as to plot the results are available at https://github.com/TBooker/Assisted-Gene-Flow and in the Dryad data repository (link to be added upon acceptance for publication). The results of all our simulations can be visualized using a Shiny App (https://shiney.zoology.ubc.ca/whitlock/AGF/), the code for which is available in the github repository.

## RESULTS

One of the goals of assisted gene flow is to provide populations with alleles that may help them cope with a changing environment. In the absence of maladaptive alleles or outbreeding depression, introducing pre-adaptive alleles always increased the fitness of recipient populations, as expected (Fig. 1A, black lines, left column). For natural systems, though, outbreeding depression and/or maladaptive alleles may be difficult to identify, which motivated us to examine a wide variety of parameter combinations. Here, we show the results from 10,000 individuals, but the results for 1,000 individuals were qualitatively similar and are available for exploration in the Shiny App. Furthermore, simulations showed little inter-replicate variation (Fig. S1), so we present our results as the means of 50 simulation replicates per parameter combination.

**Figure 1.**
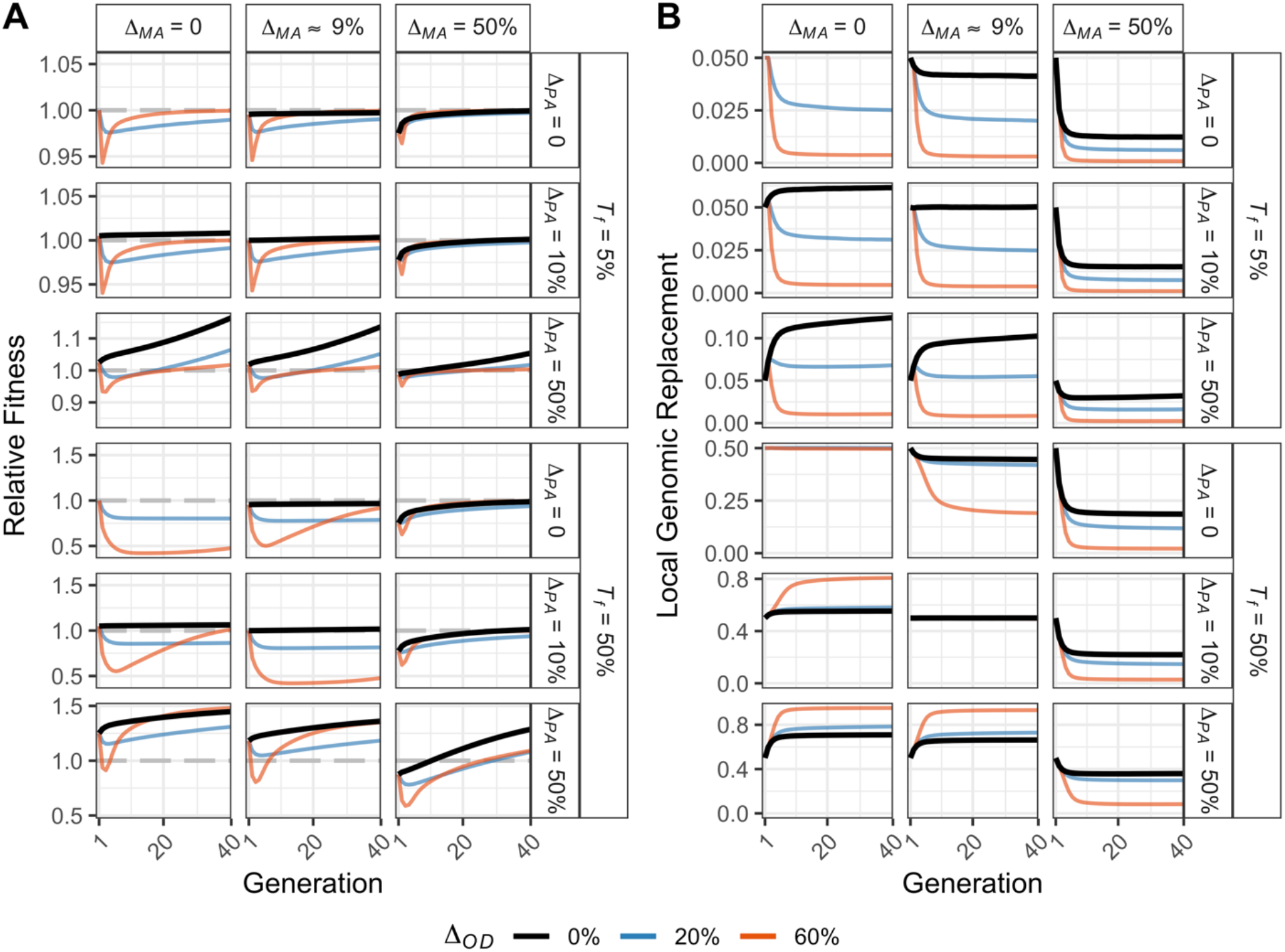
Assisted gene flow is beneficial in the long term with moderate translocation fractions (*T*_*f*_), although the mean fitness of recipient populations can be reduced for a large number of generations. (A) Relative fitness (B) and local genomic replacement fraction for the first 40 generations under *T*_*f*_ = 5% and 50% following a single translocation event in generation 1. In all cases with beneficial alleles, the mean fitness of the recipient population increased to greater than 1.0 after several generations, but the initial drop in fitness can be substantial. For this figure, genomes included ten pairs of outbreeding depression loci, and in the scenarios where pre-adaptive and maladaptive alleles occurred, five of each were present with variable selection strengths in both cases (Δ_*PA*_ and Δ_*MA*_; maladaptive with 0.5 dominance). Relative fitness value of 1.0 is indicated with a grey dashed line. Note that the scale of the *y*-axis changes for different rows and the *y*-axis limits vary in panel (B). Cases with Δ_*PA*_ = Δ_*MA*_ = 0 are not shown because they were not simulated (relative fitness would stay at 1.0).

Under a variety of outbreeding depression and maladaptation scenarios, we found that there was an initial reduction in relative fitness after a translocation replacing *T*_*f*_ = 5% of the recipient population, but the recipient population fitness typically recovered within 100 generations, and often within 20 generations (Fig. 1A upper half; Fig. S2). This was true across a range of different selection strengths and also for *T*_*f*_ = 0.5% (Fig. S3). However, with the high value *T*_*f*_ = 50%, fitness recovery was often quite delayed (Fig. 1A lower half) and did not always recover to pre-translocation levels within 100 generations when OD was > 0% and maladaptation was weak (Fig. S3). The effect of genetic incompatibilities on fitness mainly depended on the total effect across all loci rather than the number of pairs of epistatic alleles. For instance, in cases with high migration (*T*_*f*_ = 50%), scenarios with outbreeding depression of 20% frequently took longer to fully recover than with outbreeding depression of 60%, regardless of genetic architecture (Fig. 1A; Fig. S3).

The dynamics of fitness change after assisted gene flow were driven in part by the total strength of selection acting on pre-adaptive alleles (Δ_*PA*_). In the cases where the overall fitness benefit of pre-adaptive alleles was greater than or equal to the effect of maladaptation, the deleterious alleles were purged and the pre-adaptive alleles rose in frequency, increasing population mean fitness. The presence of maladaptation did not prevent pre-adaptive alleles from rising in frequency and increasing population mean fitness (Figs. 1A, S2 and S3). However, when maladaptation was stronger than the fitness benefit of the pre-adaptive alleles, the translocated individuals were purged and the assisted gene flow had negligible long-term fitness consequences for the recipient population.

The fitness consequences of the interplay between positive and negative selection were dependent on the number of individuals moved (translocation fraction). When the translocation fraction was 5% or smaller (i.e., 0.5%), a stronger selection on maladaptive alleles (Δ_*MA*_) led to slower fitness recoveries and with less ultimate increases in fitness, as mentioned above (Figs. 1A, S2 and S3). In the cases of *T*_*f*_ = 50%, increasing selection strength on pre-adaptive alleles led to fewer generations of greatly reduced fitness when selection is weak on maladaptive alleles (Δ_*MA*_ = 0).

The magnitude of population-level fitness increase due to assisted gene flow was highly dependent upon the genomic architecture underlying the traits of interest. Specifically, the strength of selection on pre-adaptive alleles had a noticeable impact on how long it took for mean fitness to increase (Fig. 2). When there was only a single large-effect pre-adaptive allele, fitness increased to the maximum possible value within ∼50 generations. However, fitness gains occurred much more slowly with genetic architectures that had more loci of weaker effect.

**Figure 2.**
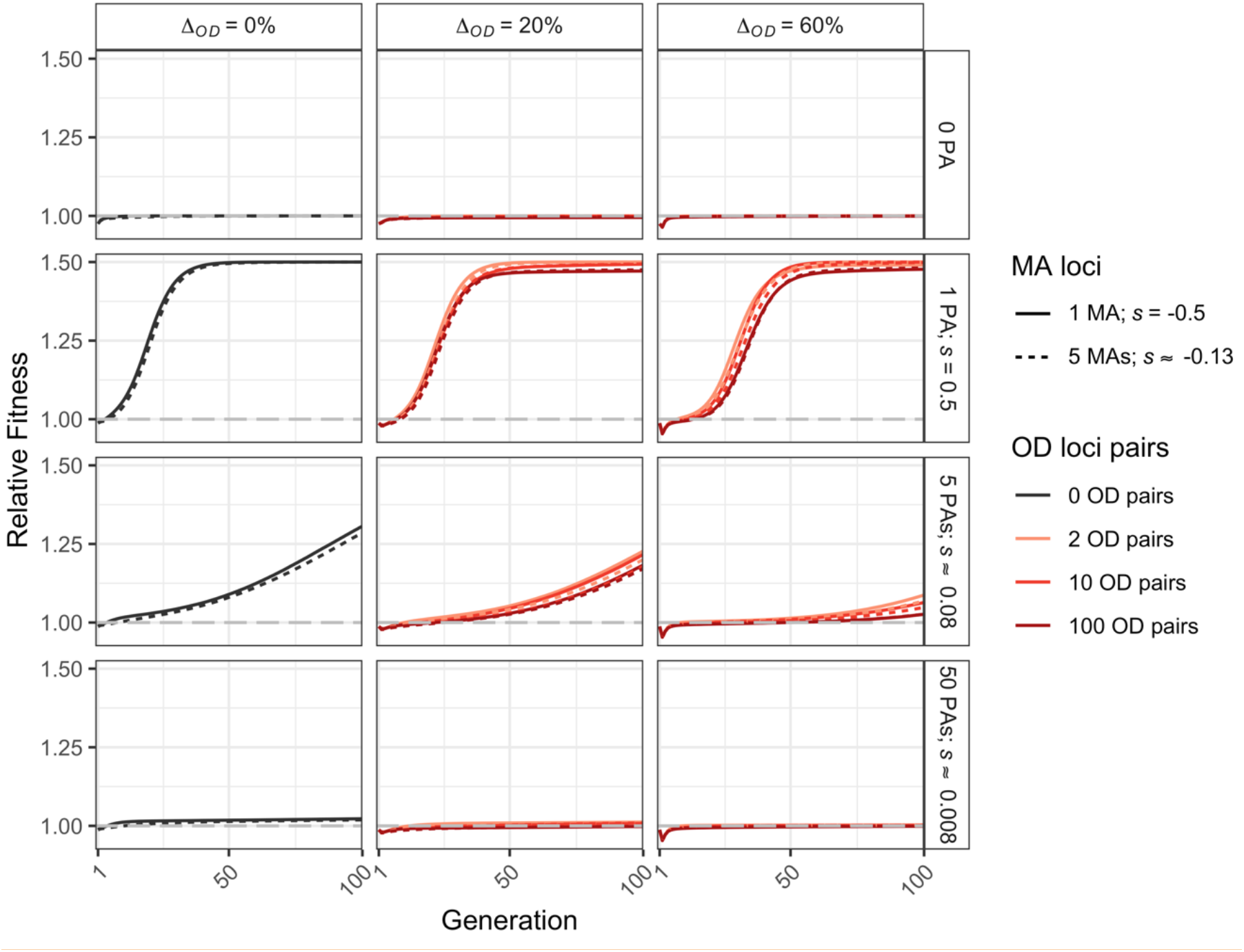
When the fitness changes of introduced alleles are the result of many loci of weak effect, the response to selection is much slower than when fewer alleles of larger effect are present. In each of these cases, the total immediate fitness effect of introduced individuals is held constant, and fitness contributions are set up as 1+ *s* per locus. Note that all plots have the same *y*-axis limits, and a relative fitness value of 1.0 is indicated with a grey dashed line. Results are shown for a single translocation event with *T*_*f*_ = 5%, Δ_*PA*_ = 50% and Δ_*MA*_ = 50% (with 0.5 dominance). Different colours denote numbers of outbreeding depression (OD) loci pairs, and the continuous and dashed lines indicate variable numbers of maladaptive alleles (MA). Note that unlike Figures 1 and 3, the *x*-axis on this plot extends to generation 100.

In contrast to the pre-adaptive alleles, the architecture of maladaptation and outbreeding depression had relatively little effect on long-term population fitness, when the translocation fraction is small (Figs. 2, S2). However, for a given strength of outbreeding depression and Δ_*MA*_, the mean fitness was slightly higher in cases with fewer outbreeding depression pairs and/or maladaptive alleles with larger effects, even though the total strength of selection was equal regardless of the number of loci (Fig. S2).

One possible concern surrounding assisted gene flow is the replacement of native genetic variation in the recipient population. We measured local genomic replacement (LGR) by calculating the proportion of donor population ancestry in the recipient population at neutral sites. We observed that local genomic replacement varies over time, but it will be less than the fraction of individuals translocated (*T*_*f*_) when maladaptation is strong (Figs. S4 and S5). When maladaptation was comparatively weak (Δ_*MA*_ < 10%), the strengths of outbreeding depression, pre-adaptive, and maladaptive alleles determined whether LGR was greater than *T*_*f*_ (Figs. 1B, S4).

The interaction of outbreeding depression and adaptation was highly dependent on the proportion of translocated individuals. For instance, when translocating a modest number of individuals (*T*_*f*_ = 5%, shown in Fig. 1B), local genomic replacement was highest when the effects of positive selection (Δ_*PA*_) outweigh negative selection (Δ_*MA*_ and Δ_*OD*_; Fig. 1B). Qualitatively similar results were obtained with a lower translocation fraction (*T*_*f*_ = 0.5%) and are presented in the Supplementary Materials (Fig. S5). When the translocation fraction was very large (*T*_*f*_ = 50%), local genomic replacement exceeded the translocation fraction value when Δ_*MA*_ was low and Δ_OD_ was high, presumably because with 50% introduction, the outbreeding depression loci are exactly at a fitness saddle and genetic hitchhiking from the PA alleles causes an increase in introduced OD alleles, leading to resolution of those loci towards introduced alleles. When the selection strengths on pre-adaptive and maladaptive alleles exactly equaled each other (Δ_*PA*_ = 10% and Δ_*MA*_ = ∼9.1%, or both equal 0.0), a stable equilibrium was reached and LGR remained at 0.5 for all outbreeding depression levels (Fig. 1B). These results were independent of dominance patterns of maladaptive loci and the number of pre-adaptive, maladaptive and outbreeding depression loci (Figs. S6, S7).

Our simulations included scenarios where the total number of introduced individuals was divided among five pulses introduced at five evenly spaced time points. We found that this pulsed migration resulted in a lower fitness reduction (e.g., relatively higher fitness) than when translocating all individuals in a single event. Indeed, we found that, when Δ_*OD*_ = 60%, the fitness reduction experienced by populations was about half that of a single translocation event (Fig. 3A); this same effect was seen with Δ_*OD*_ = 20%, but to a lesser extent (Fig. 3A). In general, the architecture of loci contributing to outbreeding depression (e.g., number of pairs of loci) had less of an impact on fitness and local genomic replacement than the overall strength of outbreeding depression (Figs. S8, S9). Additionally, with outbreeding depression, local genomic replacement (LGR) was lower in pulsed scenarios in the short-term given the smaller number of individuals introduced at each event and the frequency of introductions (Fig. 3B). It is important to note, however, that although pulsing may decrease the minimum fitness experienced (Fig. 3C), relative fitness may stay decreased for a longer period of time in comparison to when all individuals are moved at a single time (Figs. 3A, S8). After the short-term effects of pulsing have subsided, both local genomic replacement and relative fitness values of single translocation or pulsed translocation events converge on the same values after approximately 30 generations (Fig. 3A).

**Figure 3.**
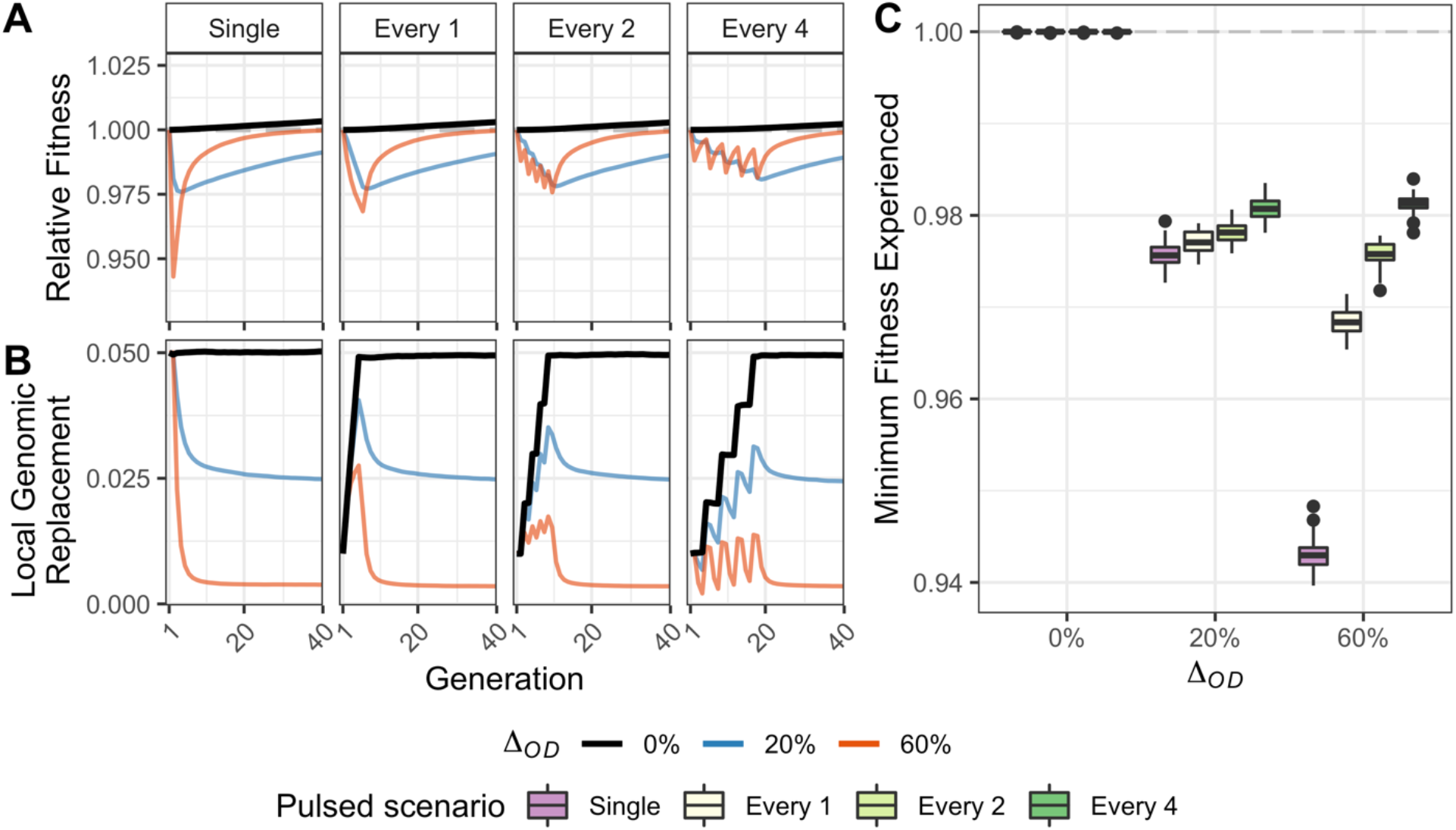
When populations suffer from outbreeding depression, pulsed introductions can decrease fitness reductions. (A) Relative fitness and (B) local genomic replacement by outbreeding depression level are shown over the first 40 generations for each translocation scenario (i.e., single translocation and five events of pulsed translocations every one, two or four generations). Note that the total number of individuals translocated was the same in each scenario (e.g., each “pulsed” introduction was one fifth the size of the single one-time introduction). Panel (C) shows the variation across 50 replicates of each translocation scenario, using the same data as in (A). Results are shown for a total migration rate of 5%, five adaptive and maladaptive alleles with a Δ_*PA*_ = 10%, Δ_*MA*_ ≈ 9.1% (dominance coefficient of 0.5), and 10 pairs of outbreeding depression loci.

In absence of outbreeding depression, introducing foreign individuals over many generations in a “pulsed” manner did not offer any measurable benefit over translocating all individuals at once (Δ_*OD*_ = 0%; Fig. 3C). Overall, changing the frequency of translocation events—whether every one, two, or four generations—offered marginal decreases in the amount of overall population fitness reduction. In these cases, considering the organismal generation time will be important— from a practical, applied perspective—in determining the frequency of translocation events. Specifically, it may be impractical to spread out translocation efforts across decades for species with long generation times.

## DISCUSSION

We have found that assisted gene flow can in some cases be a useful and powerful tool for conservation and production management. However, in many cases the advantages are small or take several generations to accrue, and the disadvantages of AGF caused by outbreeding depression and the introduction of locally maladapted alleles may have short-term consequences that need to be overcome.

### Assisted Gene Flow Leads to Modest Increases in Fitness in the Short Term

Assisted gene flow, while sometimes causing a reduction in fitness immediately following translocation, often increases population-level fitness in the long term. However, even in most beneficial scenarios, AGF does not often provide measurable benefits in the short term (e.g., the first ∼10 generations following translocation). In general, the exact fitness response resulted from a complex interplay between selection on loci with deleterious (maladaptive alleles and outbreeding depression loci) and beneficial (pre-adaptive alleles) genetic variation and their genomic architectures. The number of individual migrants (translocation fraction) had a significant impact on both the fitness response and amount of genome replaced in the recipient population. Specifically, both positive and negative fitness effects were exaggerated in the cases of higher migration levels. When the translocation effort was divided into discrete “pulses” across generations as opposed to a single translocation event, fitness reductions and genomic replacement were mitigated (particularly when outbreeding depression was strong).

Our results indicate that the conservation outcomes of AGF may be fairly modest. In our simulations, we assumed a rather extreme situation where all fitness-affecting alleles were reciprocally fixed in the donor and recipient populations. While extreme, reciprocal fixation allowed us to understand the maximum effect that AGF may have on population mean fitness. In many of the cases we tested, we found that population mean fitness had not appreciably increased even 40 generations after AGF, particularly when there were many pre-adapted loci of small effect (Figs. 1, 2, and S2A). With a translocation fraction of 5% or less, AGF was only effective when there was an oligogenic architecture of adaptation (i.e., 1 or 5 preadapted alleles) and a 50% fitness difference between the recipient and donor populations. If conservation practitioners are considering AGF as a management tool to buffer populations against the effects of anthropogenic climate change, an understanding of the genetic architecture of adaptation would be very useful. The effects of AGF on population mean fitness in long-lived species such as trees or corals, many of which are reported to have generation times in excess of decades (Babcock 1991; Howe et al. 2008), may be too slow to help populations cope with rapidly changing climates.

### The Genomic Architecture of Adaptation and Maintaining Local Identity

We examined the fitness effects resulting from different genomic architectures of three types of loci: pre-adaptive alleles, maladaptive alleles, and those causing outbreeding depression. In addition to modifying the number of these loci across the genome, we also varied their total effect. A key finding of our analysis is that the long-term fitness outcome of AGF is highly dependent upon the architecture of pre-adaptive alleles. Fitness gains were rapid when the selected trait is controlled by one or few loci of large effect. Conversely, when the trait is controlled by many loci of small effect, fitness gains were very slow and of limited benefit, even when the total possible benefit for pre-adaptive loci was the strongest (50 pre-adaptive loci and Δ_*PA*_ = 50%; Figs. 2 and S2). A similar result was found when examining the effects of genetic architecture of heat tolerance in coral, where simulated populations went extinct more quickly and had higher reductions in population size when thermal tolerance was controlled by many loci (Bay et al. 2017).

Regarding this point, it is important to understand the specific causes of fitness gains resulting from AGF. When translocating beneficial mutations only, an initial increase in fitness of the recipient population results from the translocation itself as the population now includes individuals with novel beneficial alleles. Subsequent fitness gains following this initial “fitness bump” result from selection. The change in fitness due to selection will be directly proportional to the magnitude of the selective effect on the loci controlling the trait for the following reason.

Fisher’s fundamental theorem states that the change in fitness Δ_*W*_ is equal to the additive genetic variance in fitness *V*_*A*_ (Fisher 1930; Grafen 2018). Assuming random mating, the additive genetic variance equals

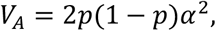

with *p* being the allele frequency, and *α* representing the slope of a regression of genotypic value on allele count (Falconer 1985; Falconer and Mackay 1996 [Fig. 7.2 therein]). Without dominance effects and with *s* being the single locus homozygous selection coefficient, this slope is 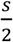, and thus

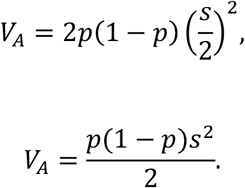

In our calculations, we held the total possible fitness benefit constant, meaning that we modelled cases with either few large-effect loci or many small-effect loci. In this case, the value of *s* decreases with the number of loci *n*_*PA*_ for a given total effect Δ_*PA*_. Although in our simulations all loci interact multiplicatively, we use the approximation of additivity in the following calculations. The approximation suffices for the illustrative purpose here. With that approximation,

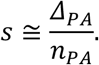

Given that we assume that the relevant alleles are reciprocally fixed in the recipient and donor populations, the initial allele frequency *p* equals the translocation fraction *T*_*f*_, and thus

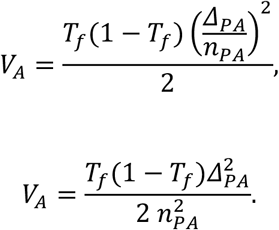

In the absence of gametic phase disequilibrium, total *V*_*A*_ is the sum of the contributions from different loci (Falconer and Mackay 1996, pg. 132):

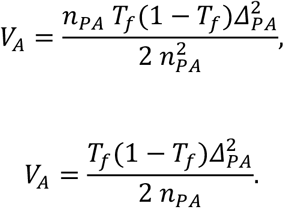

Gametic phase disequilibrium will increase additive genetic variance particularly in the early generations in our simulations (Lynch and Walsh 1998, pg. 102). However, our simulations show that this effect is not strong enough to affect the conclusion that the change in fitness decreases when an increasing number of loci are responsible for a given cumulative effect.

It follows that the change in fitness is slower for traits controlled by many loci of small effect, because additive genetic variance is smaller for these traits. An important result therefore is that for such traits, AGF is not likely to confer practically relevant benefits within a few generations for the range of parameters simulated here. On the other hand, simulations using assisted gene flow to pre-adapt *Acropora* coral to a warming climate showed that translocating as few as 10 migrants per year (with population sizes of fewer than 1,000 individuals, 114 SNPs associated with thermal tolerance, and no maladaptive loci) increased population sizes within ∼30 years, approximately 10 *Acropora* generations, of migration onset (in relation to no AGF; Bay et al. 2017).

The result that fitness gains are slow for traits controlled by many loci of small effect is an important consideration when implementing AGF, given that empirical studies have shown climate-related traits are often controlled by many small-effect loci (e.g., Rose et al. 2018; Fuller et al. 2020). For instance, drought tolerance in European populations of *Arabidopsis thaliana* was shown to be polygenic and associated with 151 SNPs (Single Nucleotide Polymorphisms; Exposito-Alonso et al. 2018). Similarly, tolerance to warm marine temperatures in *Acropora* corals was shown to be associated with variation at 114 SNPs (Bay and Palumbi, 2014). Although managers without genomic resources are at a disadvantage, it bears reminding that AGF rarely results in long-term fitness declines (Fig. S2). Furthermore, other simulations have shown little difference in the outcome of AGF when thermal tolerance is controlled by more than ca. 150 loci (Bay et al. 2017). Thus, a rough estimate of a trait’s genetic architecture or an estimation based on related species or similar traits may under some circumstances already be helpful. We also observed that the overall selection strength on maladaptive alleles (Δ_*MA*_) had a larger effect on fitness than the number of maladaptive alleles; the same pattern was seen in outbreeding depression loci (Fig. S2).

One concern that AGF raises is the loss of local genetic identity in the recipient population due to genetic swamping from the donor population, a process we term “local genomic replacement”. Through the process of AGF, some fraction of the local genome will come to be derived from outside sources, representing a change from the natural state of the population. In extreme cases, “hybridization” between donor and recipient populations could lead to the recipient population’s genomic extinction and therefore the loss of its genetic integrity (Todesco et al. 2016). Maintaining such genetic integrity (e.g., Hauskeller 2008) may be more important in some conditions (conservation) than others (optimizing resource extraction), but the case-by-case need for maintaining genetic integrity is a philosophical question beyond what we consider here (but see Rohwer and Marris 2015 for an in-depth treatment of this topic).

In the cases we simulated, the amount of local genomic replacement was largely a function of translocation fraction (*T*_*f*_). Under likely more realistic levels of translocation (*T*_*f*_ ≤ 5%), the amount of local genomic replacement was typically less than the translocation fraction. However, this replacement of local genetic variation by translocated alleles is greatest when there are the greatest fitness benefits of AGF. The exceptions were the combinations of weak total selection strength on maladaptive alleles (Δ_*MA*_ < 10%) while selection was strong on pre-adaptive alleles (Δ_*PA*_ = 50%; Fig. S5). When a large number of individuals was translocated (*T*_*f*_ = 50%), the amount of genomic turnover exceeded 80% in some cases (Fig. S5). It is important to note, however, that such a large translocation fraction is likely not realistic in a management scenario. In large populations, it is likely unfeasible to move so many individuals, and in small populations where this fraction can be achieved, inbreeding depression will likely become an issue and the results here may be inapplicable. In some real-world cases, such as reforestation of harvested sites following clearcutting, many millions of individuals may be replanted across a species range in a year. In British Columbia, for example, 259 million trees of various species were replanted in 2016 (https://news.gov.bc.ca/factsheets/factsheet-reforestation-in-bc), so a simulated translocation fraction of 50% helps illustrate fitness trends for management considerations.

### Pulsed Translocation Efforts

We examined whether dividing the translocation effort into discrete “pulses”, here represented as 20% of the total number of individuals to be moved in five separate events, had an effect on the recipient population’s fitness response. Overall, in comparison to translocating all individuals in a single effort, pulsing alleviated some of the negative fitness effects resulting from assisted gene flow. In particular, translocating individuals every four generations resulted in marked decreases in fitness reductions (e.g., relatively higher fitness) compared to more frequent pulses or a single migration (Fig. 3C). As new deleterious genotypes are introduced in each pulse, selection reduces their frequency over the subsequent generations until the next pulse of individuals arrives. This pattern, however, is largely, but not exclusively, driven by scenarios with high levels of outbreeding depression (60%). Such high levels of genomic incompatibilities via epistatic interactions are not likely to result from inter-population matings within a species; this level of outbreeding depression is more likely to result from interspecific matings and therefore not applicable to many cases where AGF may be implemented. However, the potentially unrealistic value of outbreeding depression demonstrates an extreme and shows a trend of its effect on fitness.

The genomic architecture of loci causing outbreeding depression also interacted with pulsing the translocation effort. In general, the overall strength of outbreeding depression (Δ_*OD*_) had a larger effect on the resulting fitness than the number of pairs of outbreeding depression loci (Fig. S8). Nonetheless, within a given strength of outbreeding depression, populations with fewer pairs of outbreeding depression loci experienced faster gains in fitness and higher levels of local genomic replacement (Fig. S8). This is likely because selection can more easily remove individuals from the population with fewer pairs of outbreeding depression loci, each with a higher *s*_*OD*_ value, than when many weaker pairs of outbreeding depression loci are spread across the genome and potentially linked with beneficial (e.g., pre-adaptive) alleles.

Although populations show reduced levels of both fitness reductions and local genomic replacement as a result of pulsing (Fig. 3), they also experience these depressed levels for a longer period of time. In other words, translocating individuals in a single effort might lead to lower fitness and higher local genomic replacement, but both levels recover (towards 1.0 and 0.0, respectively) more quickly. Similarly, whereas pulsing can mitigate fitness reductions when outbreeding depression is strong, it can also delay the fitness benefits of gene flow when introduced individuals have a high fitness (high Δ_*PA*_, low Δ_*MA*_) and weak outbreeding depression (low Δ_*OD*_). Hence, if pulsing helps reduce fitness reductions in some scenarios, it also delays fitness gains in other scenarios. Given that fitness and local genomic replacement levels converge on the same values regardless if pulsing was performed or not, it becomes important to consider the more critical state—a more severe fitness reduction for less time, or a less severe reduction in fitness for more time.

### Considerations for Resource Managers

We simulated a fitness increase of 50% owing to pre-adaptive alleles, meaning that individuals from the donor population would have a 50% higher fitness in the new environment. Empirical studies in natural populations have found that the strength of local adaptation is of that order in a wide variety of species (Bontrager et al. 2020). Similarly, Exposito-Alonso et al. (2019) reported strong climate-mediated natural selection in *A. thaliana* from common garden transplants where > 60% of populations were killed due to non-native (hot and dry) conditions. However, estimating the beneficial effect of pre-adaptive alleles in a novel ecological and genomic context is extremely difficult in natural settings. Furthermore, if an environment is predicted to change in a particular direction over time (e.g., trend of climatic warming), the selective benefits of pre-adaptive traits may increase in the future. In general, the benefits of pre-adaptive alleles from assisted gene flow may take many generations to realize (Figs. 1, S2), and an important consideration must be made as to whether the long-term gains outweigh the short-term fitness costs.

Assisted gene flow has been proposed as part of a decision-tree for managing coral reef restoration (Van Oppen et al. 2017). In such systems, our results can be used to guide decisions for managers, but we are aware that estimates for many of the parameters we have simulated here will not be available in most systems. It is therefore important to consider the results presented here qualitatively and in relative terms. For instance, little is known about outbreeding depression and its underlying genetic mechanisms in many systems. Our results show that outbreeding depression should be a consideration mainly when it is strong, e.g., between very divergent populations representative of interspecific crosses (Fig. S2). Given that assisted gene flow is typically done with closely related populations, outbreeding depression is not likely to play a strong role in reducing the benefits of AGF. Similarly, little is known about alleles originating in source populations that are deleterious in the recipient population. Our results highlight that this parameter (Δ_*MA*_, the maximum possible fitness reduction in an individual with all maladaptive alleles) matters when it is strong, e.g., when fitness is reduced in an individual by >10%.

This research has also generated some suggestions for managers considering assisted gene flow. First, performing controlled breeding trials before going “all-in” at the population-scale in the wild may be helpful. Many problems resulting from outbreeding depression or maladaptive alleles could be screened by measuring growth and fitness in F_1_ (and F_2_ and beyond) individuals resulting from donor-recipient crosses. Indeed, using F_1_ individuals in AGF attempts may help reduce the fitness reduction the population may experience. Second, translocating fewer individuals at a time (i.e., a smaller translocation fraction) is one way to mitigate population-level harm if breeding trials are not able to occur before translocation and unforeseen risks manifest in reductions of individual-level fitness and fecundity. Furthermore, translocating fewer individuals per translocation event mimics our pulsing scenarios that resulted in benefits previously mentioned. And lastly, if these options are not available, landscape genomic techniques that merge species distribution models with the knowledge of adaptive loci can generate recommendations for assisted migrations (Shryock et al. 2020).

In spite of a broad parameter space that we explored in our simulations, we did not examine all factors that might be considered while deciding to perform assisted gene flow or not. For instance, we did not model carrying capacity or a fluctuating population size. Secondly, we did not consider other consequences of translocations, such as disease/parasites or disruptions of social structures. Furthermore, even though adaptive genetic variation may help a population cope with environmental change, climatically-induced range shifts may increase interspecific competition in certain contexts (Razgour et al. 2019), which is a factor we did not consider.

### Future Work

Our study has provided a deepened understanding of some of the genetic factors determining the outcomes of assisted gene flow. Nonetheless, the results and parameter choices we made have exposed some avenues that future research should pursue. First, we assumed that the beneficial (pre-adaptive) alleles were reciprocally fixed between donor and recipient populations. This is likely to be an oversimplification—it is possible that these pre-adaptive alleles may already be present at a low frequency in the recipient population. Thus, what is the benefit of assisted gene flow when the pre-adaptive alleles are already present in the recipient population? Exploring the cases in which a translocation fraction (*T*_*f*_) of 0.5% was simulated may give an approximate representation of pre-adaptive alleles being present at a low frequency. In this case, however, the pre-adaptive alleles will be in linkage disequilibrium (at least in the early generations following translocation), which might not be an accurate representation of these alleles existing at low frequencies in natural populations. On a related point, further research is necessary into the uncertainty of positive selection strength of pre-adaptive loci. We modeled a constant positive selection strength over time, but selection strength may increase in the future with a changing environment. In such cases, the benefits from AGF may be more frequent or strong than our results imply.

Although we were primarily concerned with assisted gene flow as a means to improve the overall genetic health and productivity of a population and therefore measured relative fitness, it would also be worthwhile to explore the impacts of assisted gene flow on population size. Given that increasing population size is one of the principal goals of genetic rescue, population size is perhaps more comprehensible than fitness and almost certainly to be of interest to resource managers. Future research might consider allowing population size to fluctuate instead of maintaining a fixed size, and might explicitly model the possibility of population extinction.

Lastly, much remains to be explored in terms of interactions between divergent genomes and the effects of outbreeding. Although we modeled such genomic interactions as only having zero or negative consequences (e.g., outbreeding depression), we did not explore how “hybrids” between donor and recipient individuals may have hybrid vigor (e.g., heterosis). Although relatively little is known about heterosis in natural populations, it is expected to be strongest in small populations (Whitlock 2002). Although our study was concerned with large populations, the use of translocations to promote genetic rescue in small populations has recently received increasing attention (Bell et al. 2020).

## Conclusions

As climate change intensifies and populations experience fitness reductions and/or local extinctions, management strategies such as assisted gene flow will become a more widely considered tool for “prescriptive evolution” (Smith et al. 2014). Our results indicate that the conservation outcomes of AGF may be fairly modest in real world settings. In our simulations, we assumed a rather extreme situation where all fitness-affecting alleles were reciprocally fixed in the donor and recipient populations; this allowed us to understand the maximum effect that AGF may have on population mean fitness. In many of the cases we tested, we found that population mean fitness had not appreciably increased even 40 generations after AGF, particularly when there were many pre-adapted loci of small effect (Figs 1, 2, and S2A). If the alleles that contribute to local adaptation have individually weak fitness effects, the effects of AGF on population mean fitness in long-lived species may be too slow to help populations cope with rapidly changing climates. With a translocation fraction of 5% or less, our simulations showed AGF to be effective only with a 50% fitness difference between the recipient and donor populations and when an oligogenic architecture of adaptation underlied the adaptive trait (i.e., 1 or 5 preadapted alleles) Fig. 2).

Although detailed knowledge of outbreeding depression and the genetic architecture of adaptive (both pre- and maladaptive) traits would greatly improve predictions regarding the long-term success of assisted gene flow, such knowledge is often rudimentary at best, and often limited to model systems. However, tools such as controlled breeding trials or landscape genomics can help inform managers before conducting AGF. Although assisted gene flow has the potential to lead to negative short-term fitness consequences, its long-term benefits suggest it may be a useful management tool moving forward to help populations adapt to a changing climate.

## Acknowledgements

Thanks to Sally Aitken for discussion and thanks to Sam Yeaman and Moisés Expósito-Alonso for comments on the manuscript. Additionally, comments from UBC Biodiversity Research Centre participants of an early presentation of this work increased its quality and clarity.

Funding for parts of this study was provided by a Genome Canada Large-Scale Applied Research Project in Natural Resources and the Environment (Project code 242RTE) to JAG; TRB was funded by the CoAdapTree project which is funded by Genome Canada (241REF), Genome BC and 16 other sponsors (http://coadaptree.forestry.ubc.ca/sponsors/); the Swiss National Science Foundation (SNF) Doc. Mobility fellowship to RMD (P1SKP3_168393); the Swiss National Science Foundation to PN (P400PB_180870); a BRITE postdoctoral fellowship to ATT from the Biodiversity Research Centre at the University of British Columbia; and an NSERC Discovery Grant to MCW.

## Author Contributions

MCW conceived of the study. RMD performed genetic simulations, TB analyzed the data and AT and TB made the figures. JAG led the manuscript writing. All authors designed the study and contributed to manuscript writing and editing.

