## Supplemental Figures for "The genetics of assisted gene flow: immediate costs and long-term benefits"

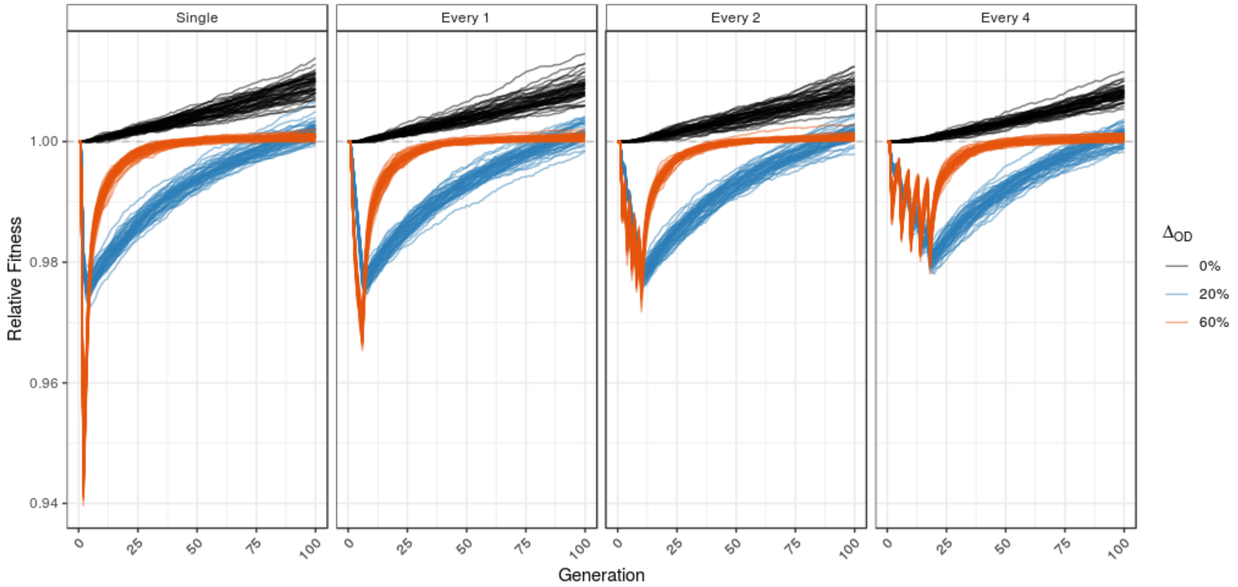


**Supplemental Figure S1**. Variation among simulation replicates within the same parameter combination: 10,000 individuals, translocation fraction ($T_f$) = 5%, 10 outbreeding depression loci, 5 pre-adaptive and 5 maladaptive loci, selection strenths $\Delta_{PA}$ = 10% and $\Delta_{MA}$ = ~9%, and a maladaptive allele dominance coefficient of 0.5.

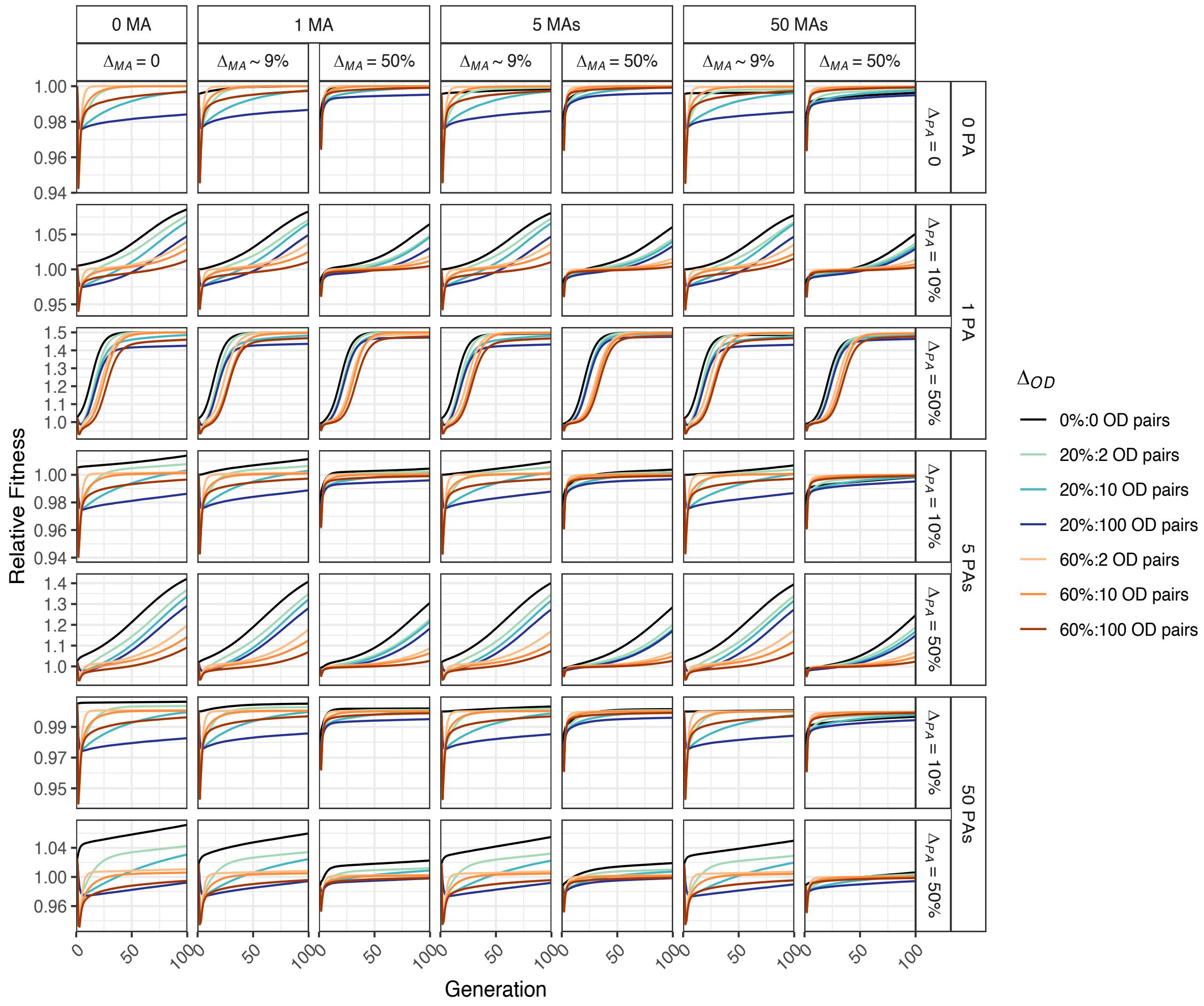


**Supplemental Figure S2**. Relative fitness when varying selection strength ($\Delta_{PA}$, $\Delta_{MA}$, $\Delta_{OD}$) and number of pre-adaptive (PA) alleles, maladaptive (MA) alleles, and outbreeding depression (OD) loci. These results use the same parameters as Figure 1 in the main text (including a single translocation event with a translocation fraction of 5%).

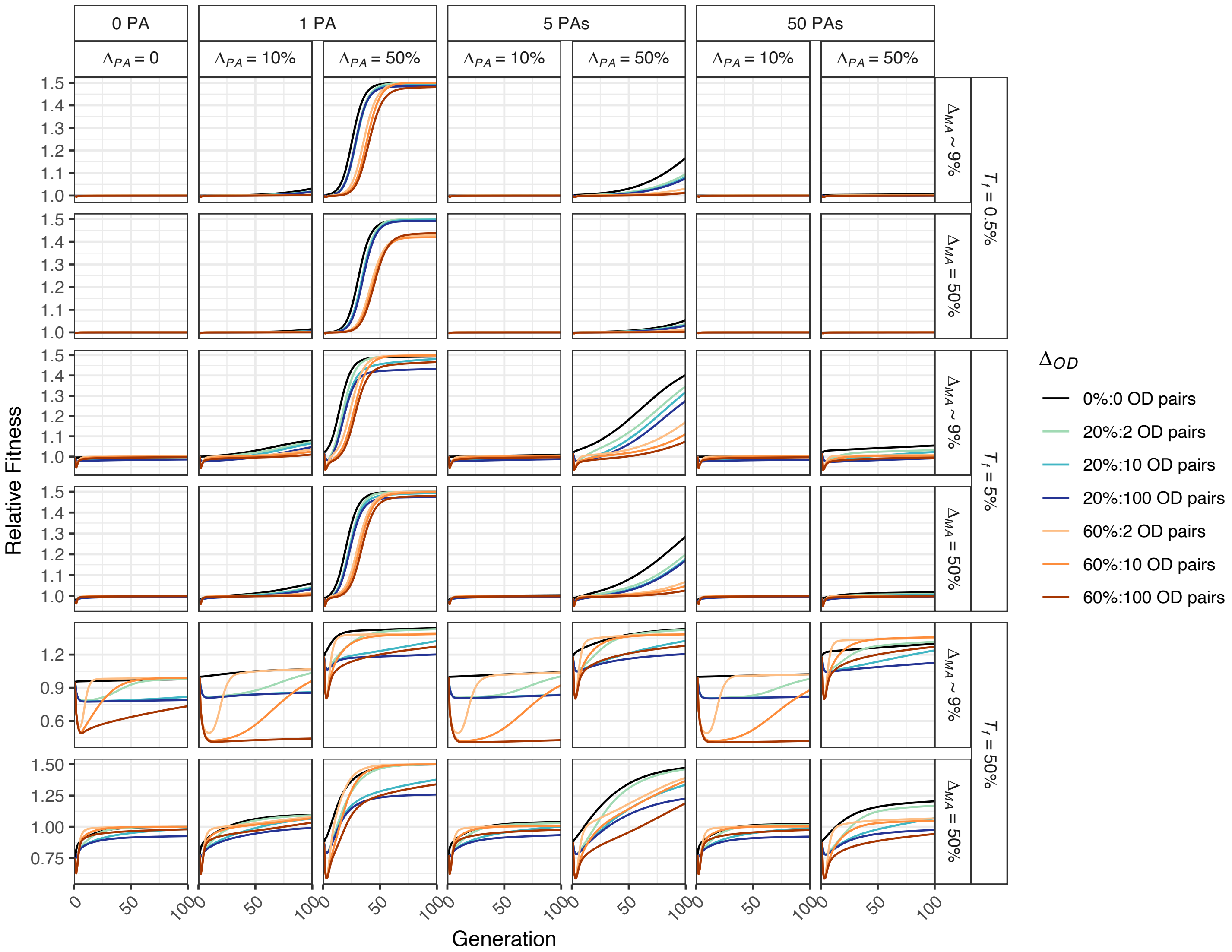


**Supplemental Figure S3.** Relative fitness with five maladaptive loci and a single translocation event ($T_f$) of 0.5%, 5%, or 50%. Selection strength is variable for pre-adaptive alleles ($\Delta_{PA}$), maladaptive alleles ($\Delta_{MA}$), and outbreeding depression alleles ($\Delta_{OD}$), as were the numbers of pre-adaptive (PA) and outbreeding (OD) loci; maladaptive allele dominance coefficient of 0.5.

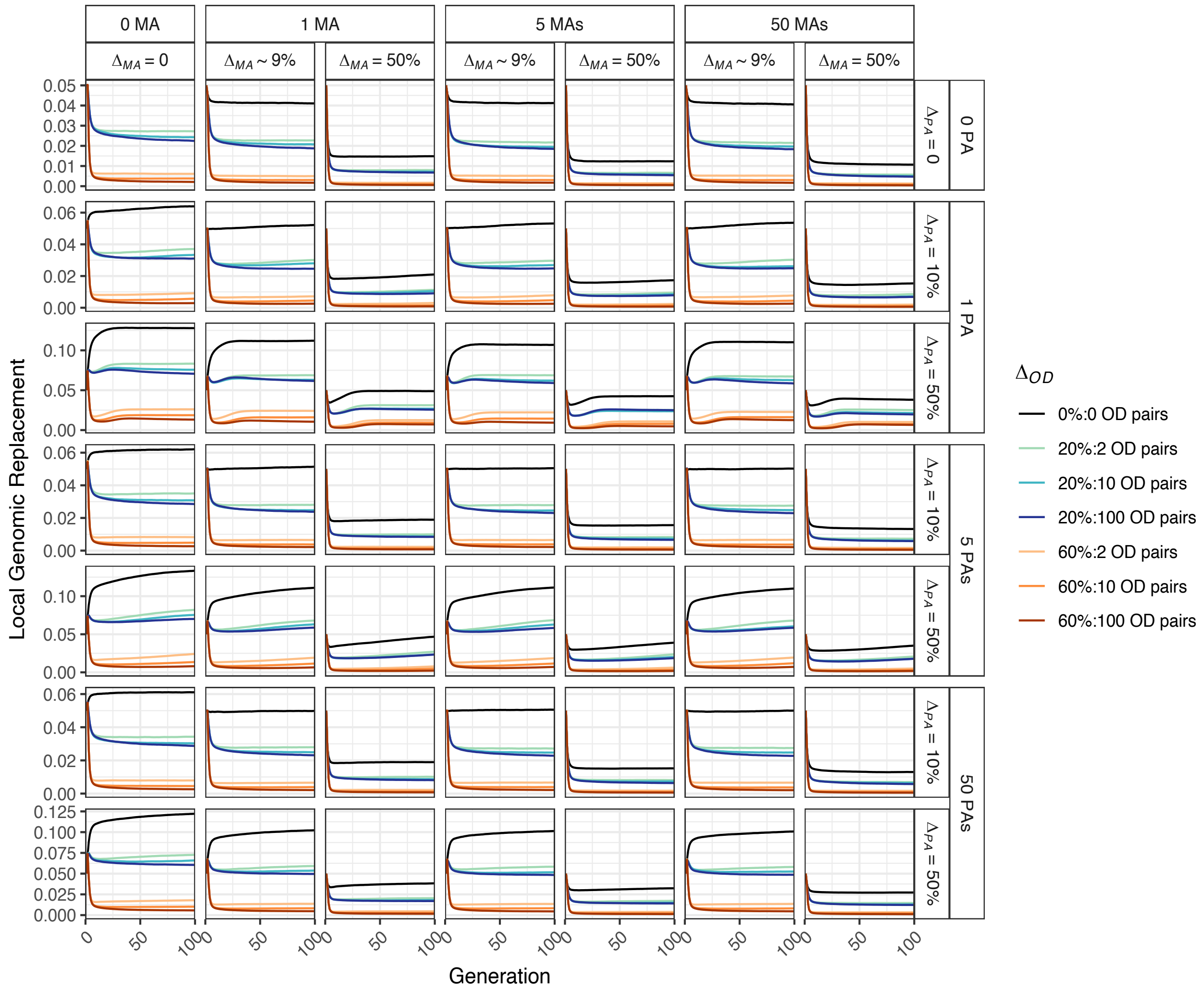


**Supplemental Figure S4**. Local genomic replacement when varying selection strength ($\Delta_{PA}$, $\Delta_{MA}$, $\Delta_{OD}$) and number of pre-adaptive (PA) alleles, maladaptive (MA) alleles, and outbreeding depression (OD) loci. These results use the same parameters as Figure 1 in the main text (including a single translocation event with a translocation fraction of 5%).

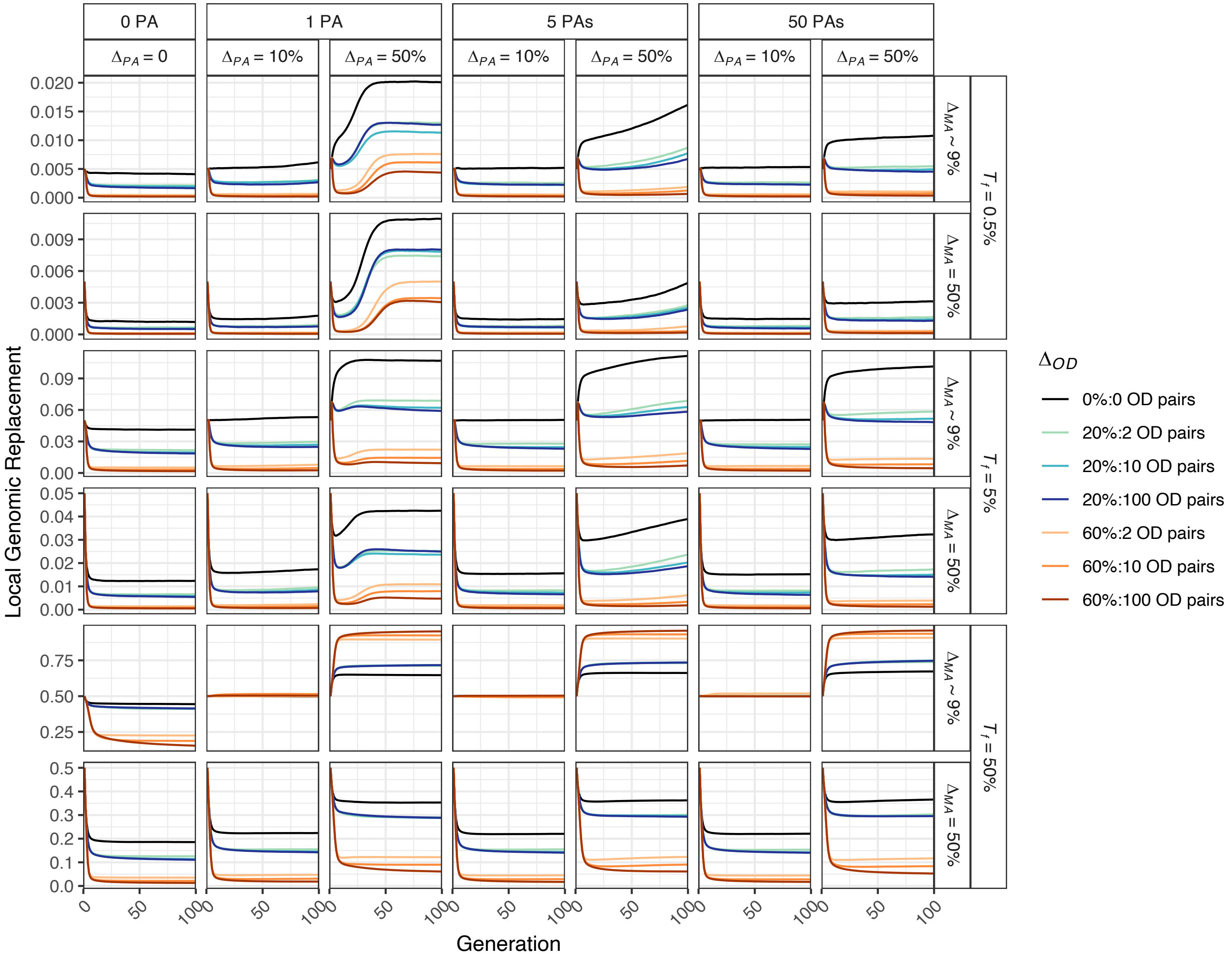


**Supplemental Figure S5.** Local genomic replacement (B) with five maladaptive loci and a single translocation event ($T_f$) of 0.5%, 5%, or 50%. Selection strength is variable for pre-adaptive alleles ($\Delta_{PA}$), maladaptive alleles ($\Delta_{MA}$), and outbreeding depression alleles ($\Delta_{OD}$), as were the numbers of pre-adaptive (PA) and outbreeding (OD) loci; maladaptive allele dominance coefficient of 0.5.

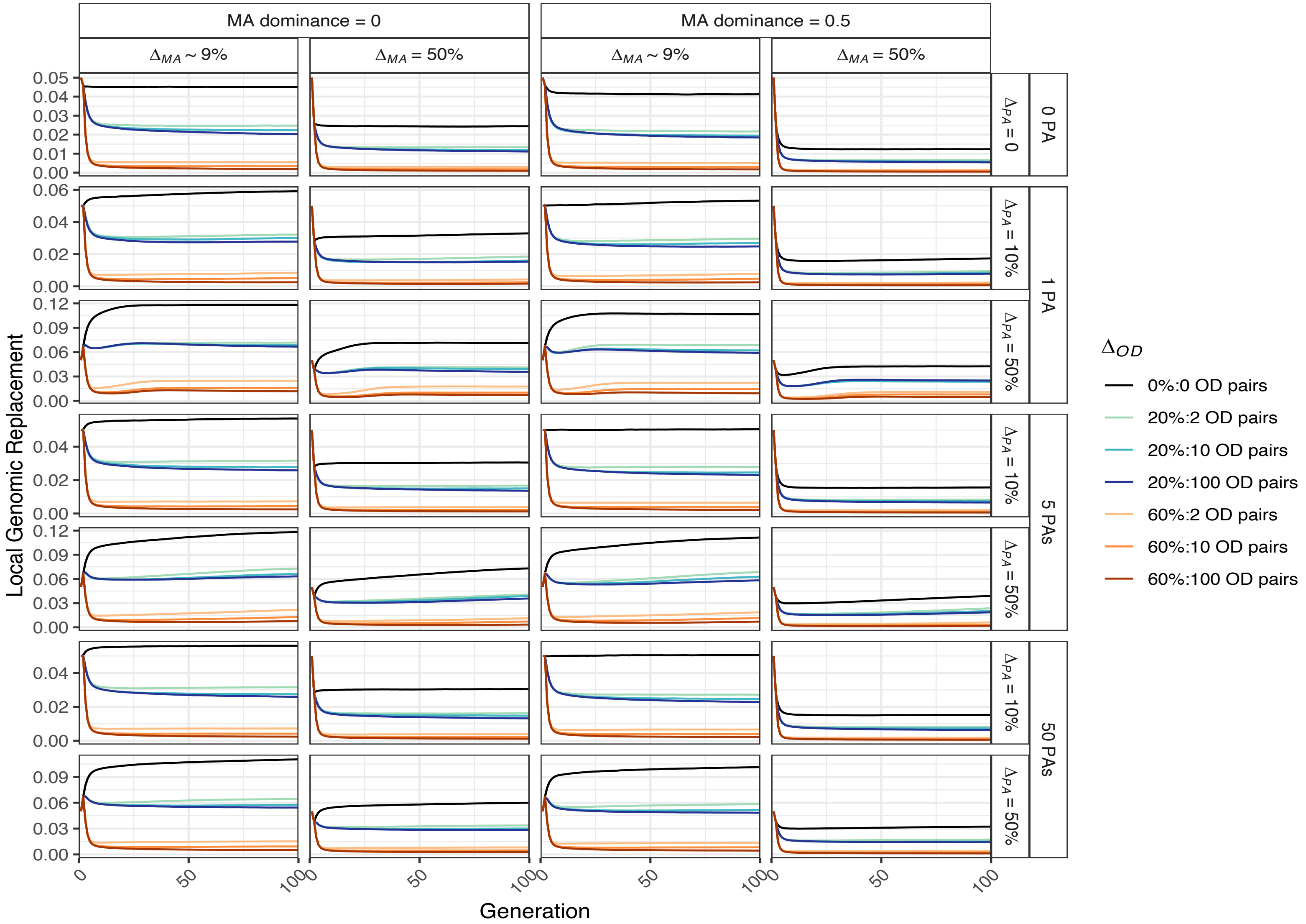


**Supplemental Figure S6.** Local genomic replacement as a function of maladaptive allele (MA) dominance and a single translocation event ($T_f$) of 5%. Selection strength is variable for pre-adaptive alleles ($\Delta_{PA}$), maladaptive alleles ($\Delta_{MA}$), outbreeding depression alleles ($\Delta_{OD}$), and the numbers of pre-adaptive (PA) and outbreeding (OD) loci, but the number of maladaptive loci was fixed to 5.

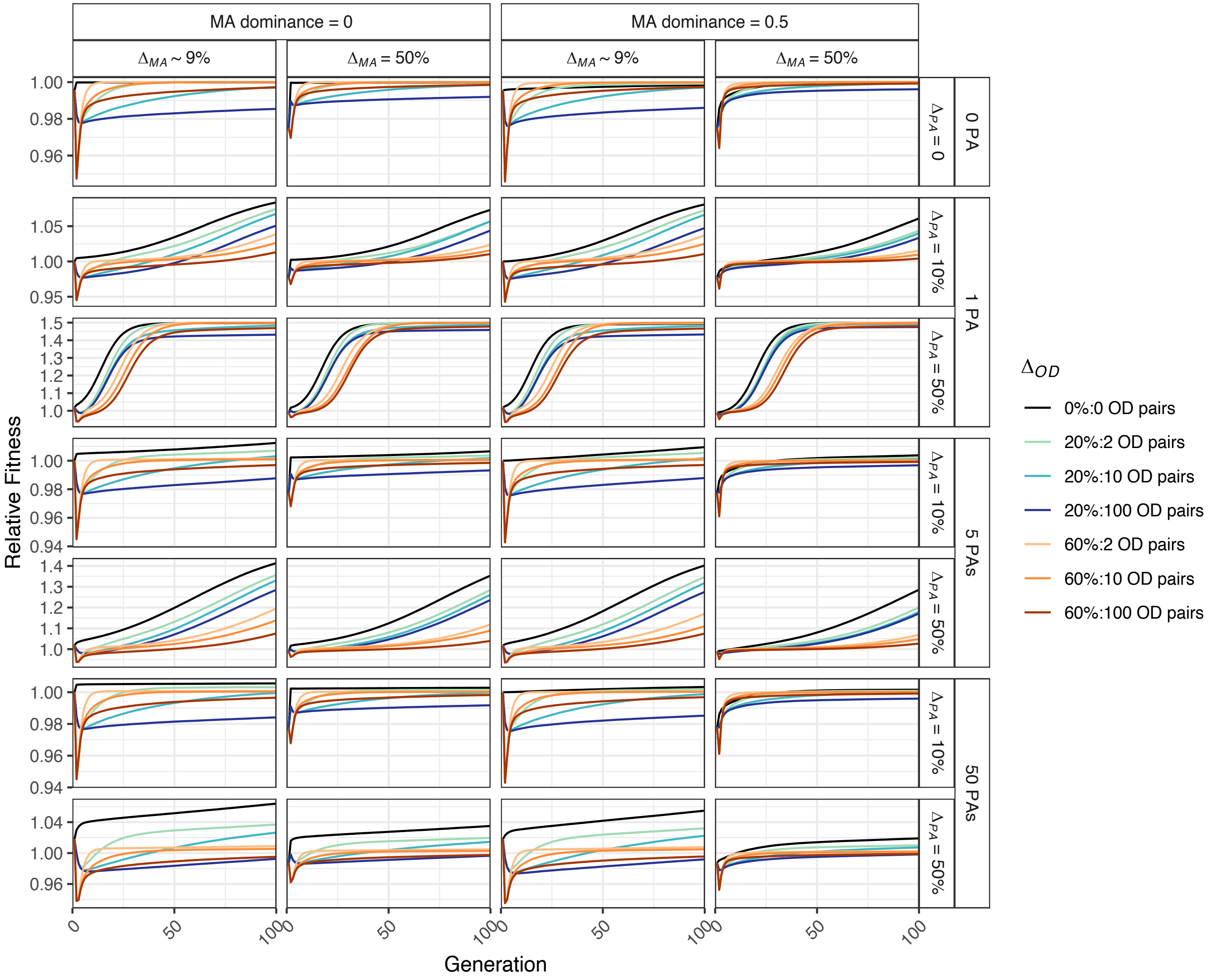


**Supplemental Figure S7.** Relative fitness as a function of maladaptive allele (MA) dominance and a single translocation event ($T_f$) of 5%. Selection strength is variable for pre-adaptive alleles ($\Delta_{PA}$), maladaptive alleles ($\Delta_{MA}$), outbreeding depression alleles ($\Delta_{OD}$), and the numbers of pre-adaptive (PA) and outbreeding (OD) loci, but the number of maladaptive loci was fixed to 5.

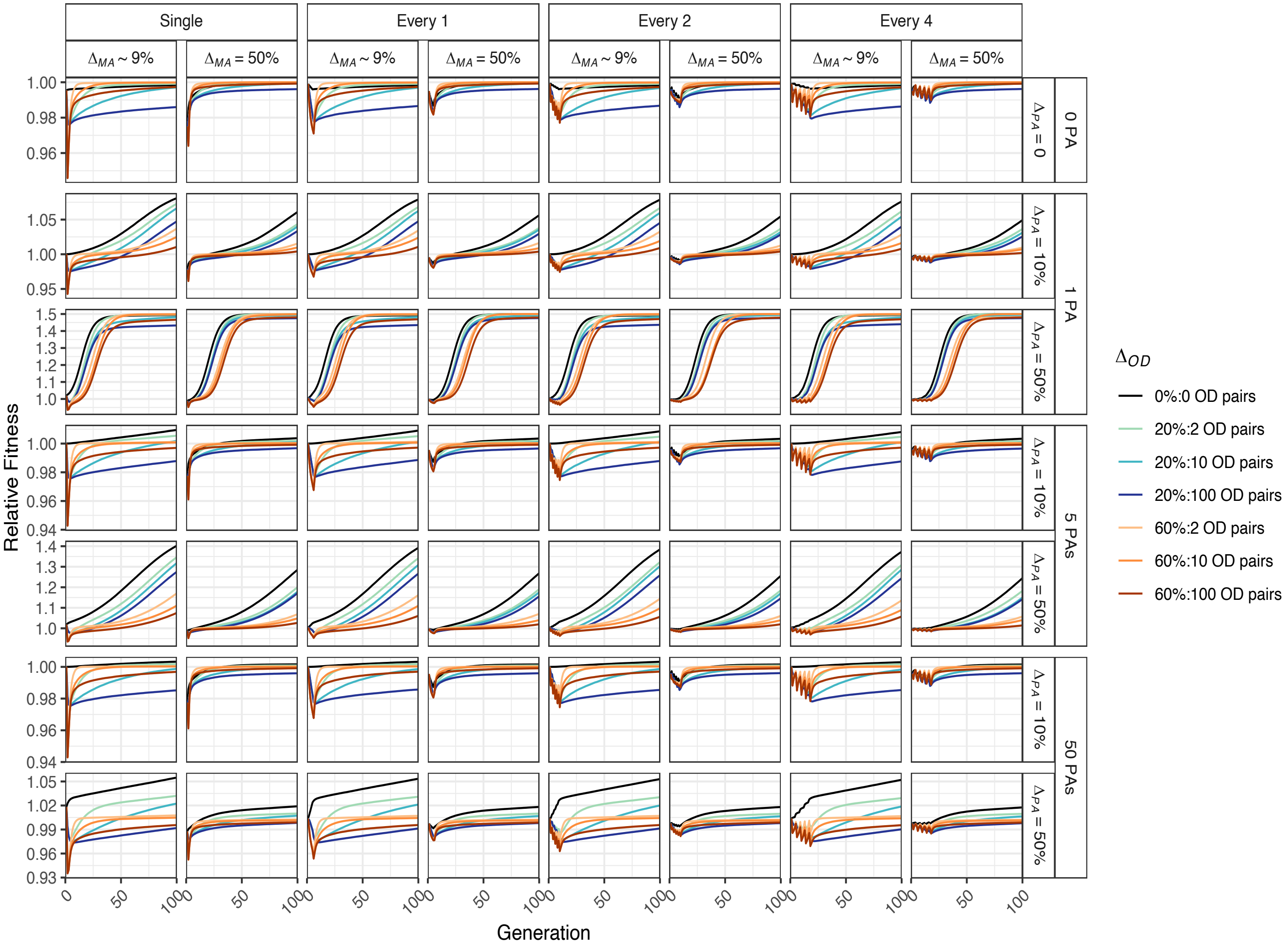


**Supplemental Figure S8.** Relative fitness when comparing a single translocation event of 5% ($T_f$) to a "pulsed" translocation where 20% of the translocated individuals were introduced in each of five generations separated by one, two, or four generations. Selection strength is variable for pre-adaptive alleles ($\Delta_{PA}$), maladaptive alleles, ($\Delta_{MA}$) and outbreeding depression alleles ($\Delta_{OD}$), as were the numbers of pre-adaptive (PA) and outbreeding (OD) loci. Five maladaptive loci are present, and maladaptive allele dominance equals 0.5.

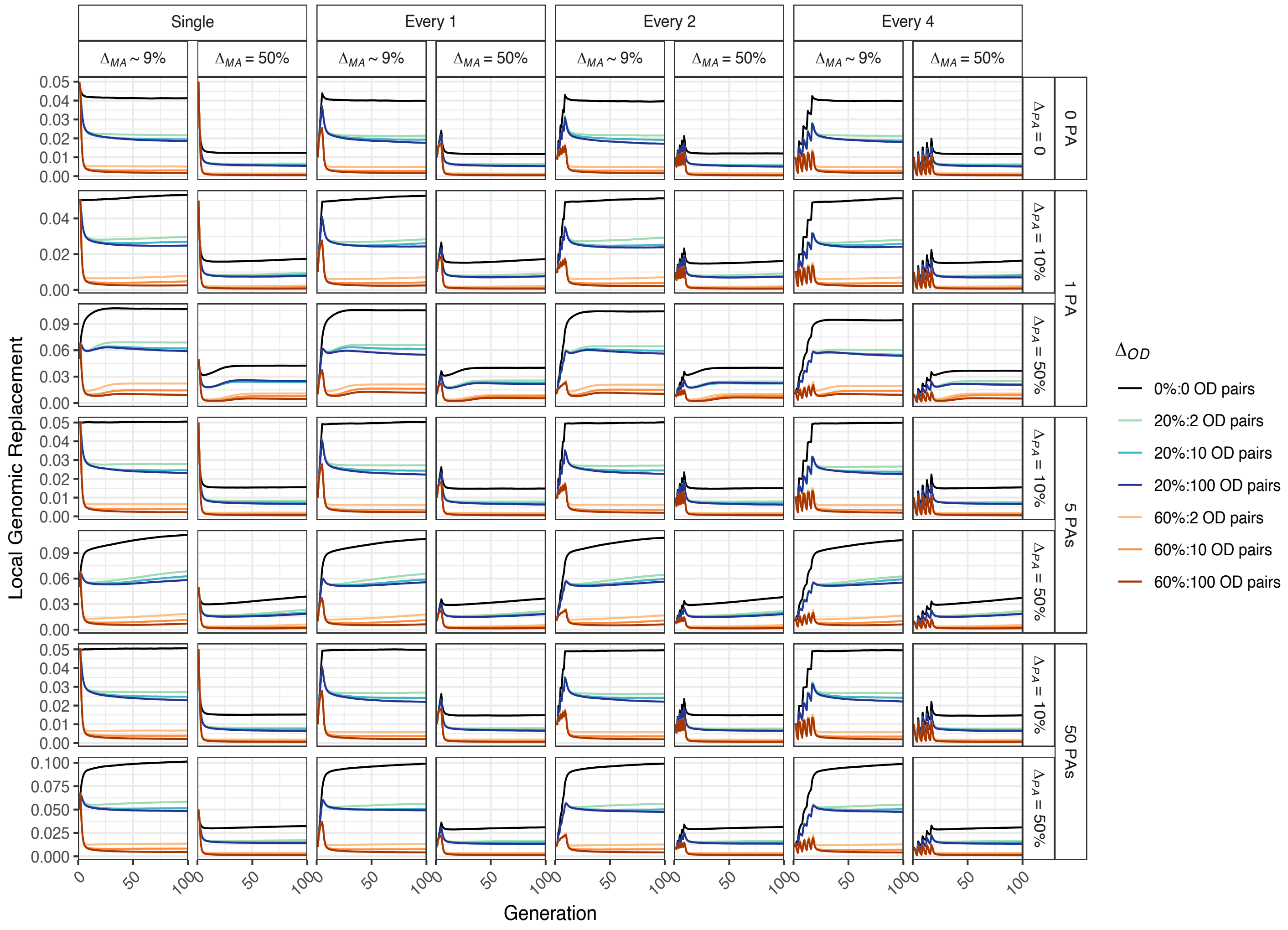


**Supplemental Figure S9.** Local genomic replacement when comparing a single translocation event of 5% ($T_f$) to a "pulsed" translocation where 20% of the translocated individuals were introduced in each of five generations separated by one, two, or four generations. Selection strength is variable for pre-adaptive alleles ($\Delta_{PA}$), maladaptive alleles, ($\Delta_{MA}$) and outbreeding depression alleles ($\Delta_{OD}$), as were the numbers of pre-adaptive (PA) and outbreeding (OD) loci. Five maladaptive loci are present, and maladaptive allele dominance equals 0.5.
